# Nutrient enrichment intensifies plant competitive effects, favouring monocultures: a global meta-analysis

**DOI:** 10.1101/2025.09.26.678566

**Authors:** Laís Petri, Ashwini Ramesh, Alejandra Martínez-Blancas, Amar Deep Tiwari, Patrick S Bills, Lauren L. Sullivan, Phoebe L. Zarnetske

## Abstract

Excess nutrient deposition is a major driver of plant diversity loss, but the mechanisms of this loss remain unclear. Diversity loss has been interpreted through the niche dimension hypothesis, yet the mechanistic link via competitive effects remains unresolved. Establishing this link demands a global synthesis of how nutrients alter competitive effects, defined as a species’ ability to perform better or worse with heterospecific competitors than with conspecifics. Strong competitive effects from heterospecifics relative to conspecifics can lead to decline or exclusion of the focal species, whereas weaker competitive effects indicate niche differentiation that promotes coexistence. We conducted a global meta-analysis of 71 plant competition experiments to quantify how nutrient addition alters competitive effects, using biomass as the performance metric. Nutrient addition intensified competitive effects with heterospecifics by 1.5-fold, as focal species biomass was higher with conspecifics. Responses depended on the initial competitive effect under low nutrients: when effects were weak, nutrient addition increased biomass with conspecifics; when effects were strong, nutrient addition increased biomass with heterospecifics, alleviating competition and potentially countering species loss. Competitive effects were amplified most when higher temperatures occurred in dry quarters, suggesting nutrient additions may exacerbate competition under extreme climate conditions. Understanding these interactive effects of nutrient addition, changing climate, and competition is critical for predicting plant responses to global change in both managed and natural ecosystems.

**Significance Statement:** Nutrient enrichment from agricultural runoff is a driver of plant diversity loss. While patterns of diversity loss from nutrient additions are clear, the processes driving competitive effects that lead to diversity loss remain uncertain. Our global meta-analysis reveals that multiple nutrient addition intensifies competitive effects favouring monocultures i.e, species accumulating more biomass with conspecifics than heterospecifics. Depending on the initial strength of competitive effects, nutrient addition amplified or alleviated them. Temperature and water availability interact with nutrient addition to amplify competitive asymmetries, likely further shaping species dominance under changing climate. These insights provide the first global-scale evidence of nutrient enrichment effects on plant communities and highlight the need to mitigate compounding impacts of fertilization and climate change on diversity.

## Introduction

Nutrient deposition in terrestrial ecosystems results in dramatic losses in species richness (1), with up to a 35% reduction in plant species in temperate regions (2). Specifically, nutrient addition results in the loss of nitrogen-fixing plants (3, 4) and forbs (5), leading to significant and repeatable loss of plant diversity (6). The niche dimension hypothesis offers a mechanistic explanation for how competition for added nutrients leads to species loss. When species are limited by different resources, competitive trade-offs in resource acquisition allow them to coexist (7, 8). Since plant species compete for a similar set of limiting resources (e.g., soil nutrients, light, water), the addition of a limiting resource eliminates such potential trade-offs, reducing the number of species that can coexist (8). While the patterns of diversity loss under nutrient excess are clear (9–11), the mechanisms, particularly the role of competitive effects in driving shifts in species’ composition, remain elusive.

The direct manifestation of the niche dimension trade-offs can be captured by species’ competitive effects. Competitive effects, defined as a species’ ability to perform better with its heterospecific than conspecific competitors (e.g., mixtures vs. monocultures) typically quantified via biomass, yield, or relative growth rate (12, 13) (Box Fig. 1). Strong competitive effects from heterospecifics lead to the decline or exclusion of the focal species, whereas weaker effects can signal niche differentiation, a key ingredient for coexistence (14). Three decades of nutrient addition experiments have emphasized pairwise interactions as the building blocks for understanding competitive effects in diverse ecosystems (15, 16). These experiments have resulted in a general understanding that nutrient addition intensifies competitive exclusion by enhancing the dominance of species with rapid growth or efficient light capture, while disadvantaging those reliant on low-resource persistence strategies (7, 17, 18). Nutrient addition can also reduce the importance of belowground competition relative to aboveground competition (19, 20). Despite such important insights, most work has remained system specific. As a result, we still lack a synthesis that situates competitive effects within the niche dimension hypothesis, a unifying principle for how resource trade-offs could structure diversity. Therefore, the question remains: how does multiple nutrient addition shape species’ competitive effects?

**Box Fig 1.**
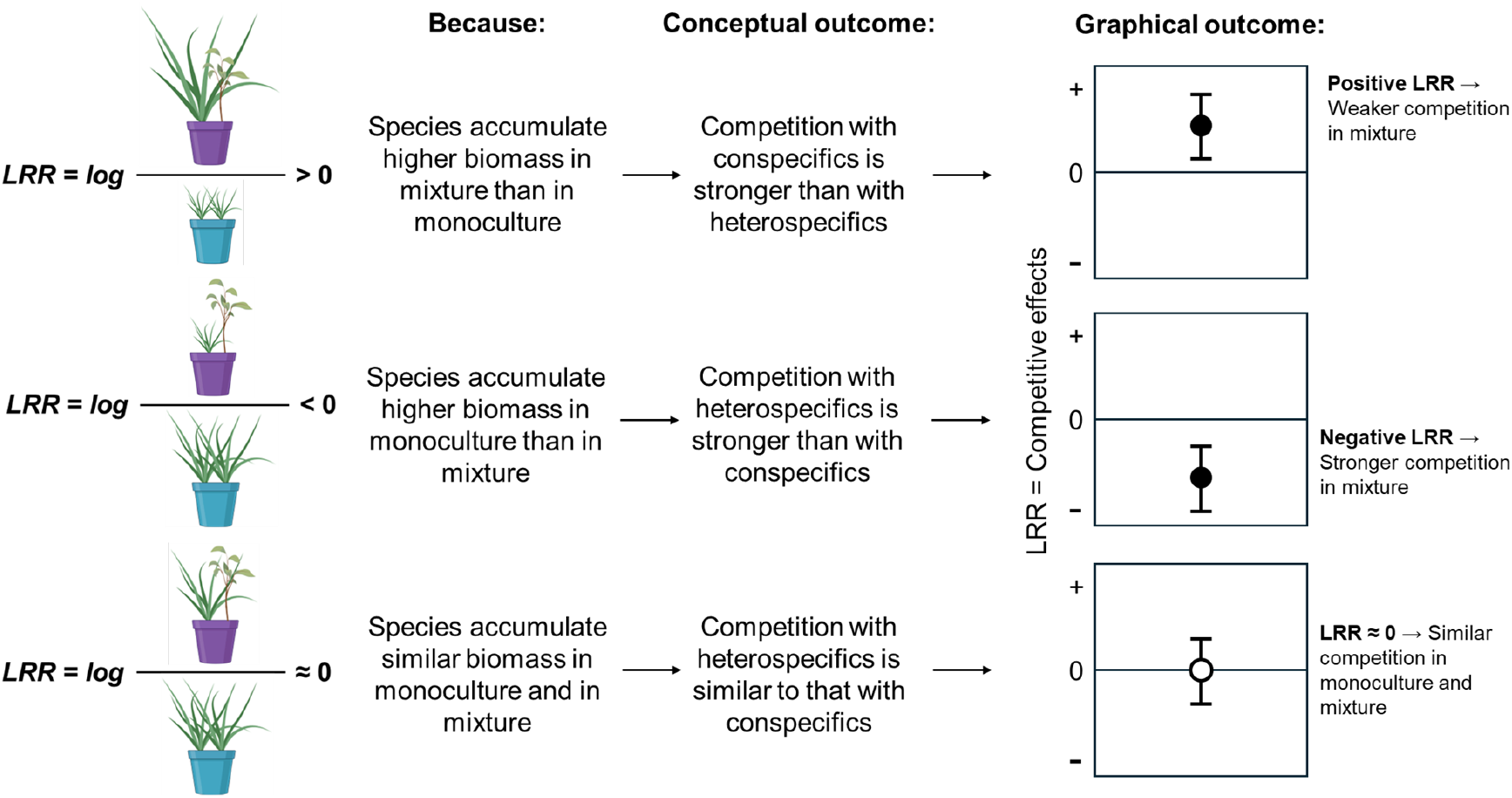
How competition experiments translate to Log Response Ratio (LRR). We used commonly performed competition treatments of species in competition with heterospecifics (purple pot) or in competition with conspecifics (blue pot) to analyse competitive effects. Plant size reflects biomass accumulation, with larger plants indicating greater biomass and smaller plants indicating less. Stronger competitive effects reduce biomass accumulation. Positive LRR values indicate stronger competitive effects from conspecifics than heterospecifics on the focal species, while negative LRR values indicate the opposite. Pots with plants created with BioRender.com.

Efforts to assess the impact of nutrient enrichment on plant competitive effects are fraught with challenges. First, nutrient addition yields inconsistent competitive effect responses (21). Added nutrients pull several competitive levers at once such that species’ competitive effects among heterospecifics may strengthen, weaken, or remain unchanged (22). Most studies do not disentangle whether divergent competitive effects stem from the amount of added nutrients (fold change) or from qualitative differences across nutrient types (e.g., N vs. NPK) (21, 23). As a result, the net outcome of nutrient addition on species competitive effects has limited predictability across ecosystems. Second, most nutrient experiments frequently rely on a small number of binary or categorical nutrient treatments rather than continuous addition gradients, limiting our ability to observe how competitive effects shift along nutrient gradients or to detect observation of nonlinear effects that shape competitive dynamics (24). Third, species interactions are often summarized as mean responses across pairs, masking the heterogeneity in competitive effects that underpins coexistence (25) (Box Fig. 2). Such averaging obscures whether some species experience strong exclusion while others retain opportunities for niche differentiation with nutrient addition. Capturing this variation requires species-level performance data across finely resolved nutrient gradients. Finally, nutrient addition via fertilization or deposition rarely occurs in isolation, but often coincides with other global change factors such as warming and drought (26). Under such combined pressures, the stress-gradient hypothesis predicts that abiotic stress can reduce competitive effects (27, 28), yet the interplay between nutrient addition and other stressors is still poorly resolved. With simultaneous warming, water scarcity, and nutrient addition on the rise, separating multiple global change pressures is crucial for forecasting shifts in competitive effects (29). Together, these gaps highlight the need for a global synthesis evaluating plant competitive ability with nutrient addition across ecosystems.

**Box Fig 2.**
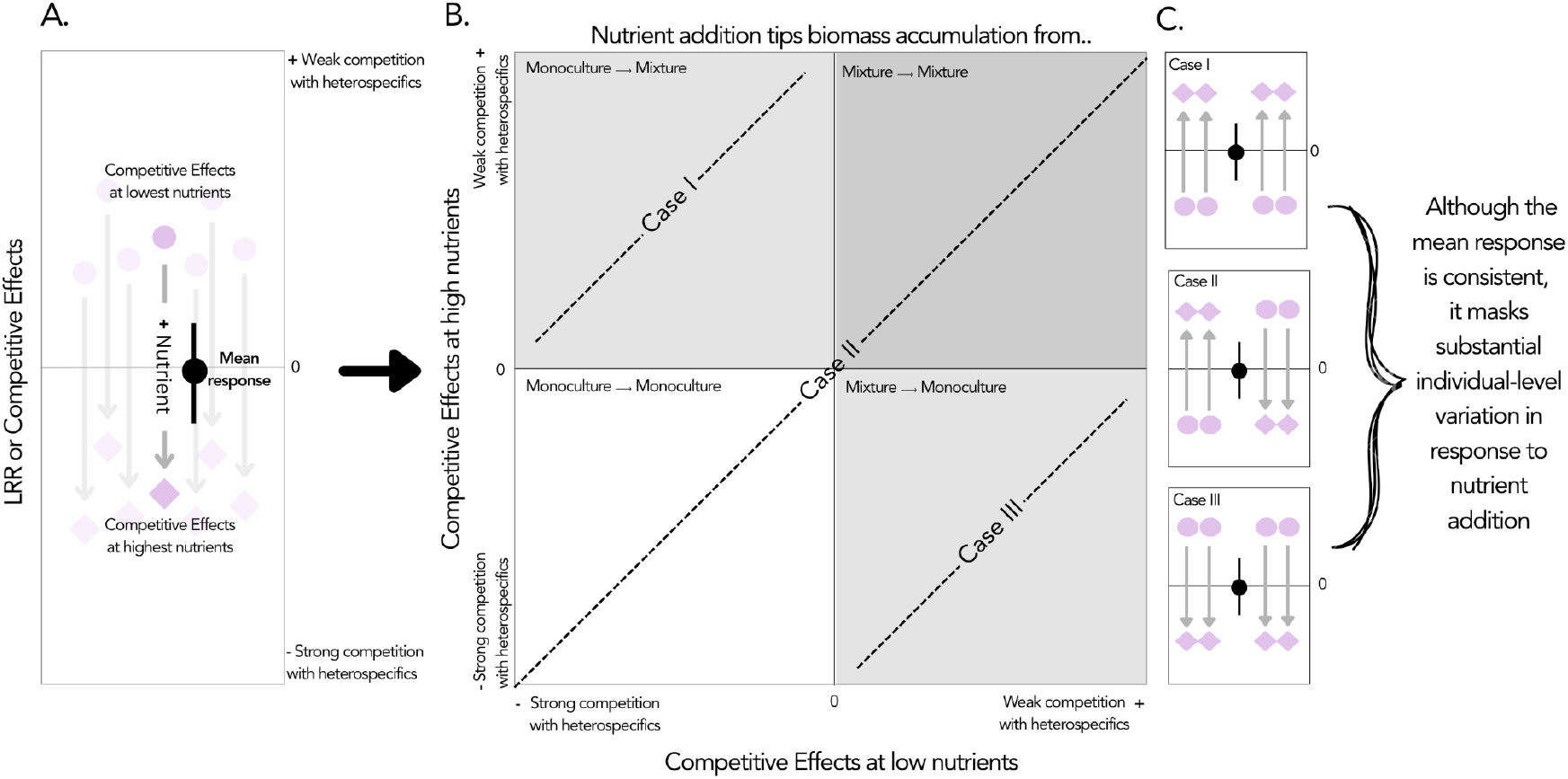
Conceptual framework showing how species’ competitive effects as measured from their log response ratio shift from low (●) to high (♦) nutrient conditions (A) can be mapped into a (B) two-dimensional space: low-nutrient competitive effects on the x-axis and high-nutrient competitive effects on the y-axis. (C) Quadrant colors represent combinations of strong or weak competitive effects under low and high nutrients. Arrows and side panels illustrate example cases of individual variation and overall mean shifts. Cases I–III show different individual responses that result in no net change in mean competitive effects.

To overcome these challenges, we performed a global meta-analysis of pairwise plant competition studies to quantify how nutrient addition affects species’ competitive effects. From over 3,500 screened papers, we extracted data from 31 studies that experimentally manipulated nutrients, compiling a species-level database to capture patterns and context-dependent responses beyond the reach of single experiments. First, we assess whether the addition of multiple limiting soil nutrients amplifies competitive effects of a focal plant growing with heterospecifics *vs*. conspecifics and by quantifying how competition strength scales with the amount of added nutrients (Q1; Figs. 1-2). Second, we evaluate variation in species response to nutrient addition, asking whether nutrient addition causes competitive effects with heterospecifics to shift at low versus highest nutrient addition (Q2; Fig. 3, Box Fig. 2). Finally, we examine how climate, particularly temperature and precipitation, modulates nutrient-driven competition (Q3, Fig. 4). Together, these analyses provide a global assessment of how nutrient addition alters competitive effects, as a necessary step toward anticipating plant diversity loss under accelerating nutrient and global change pressures.

**Fig 1.**
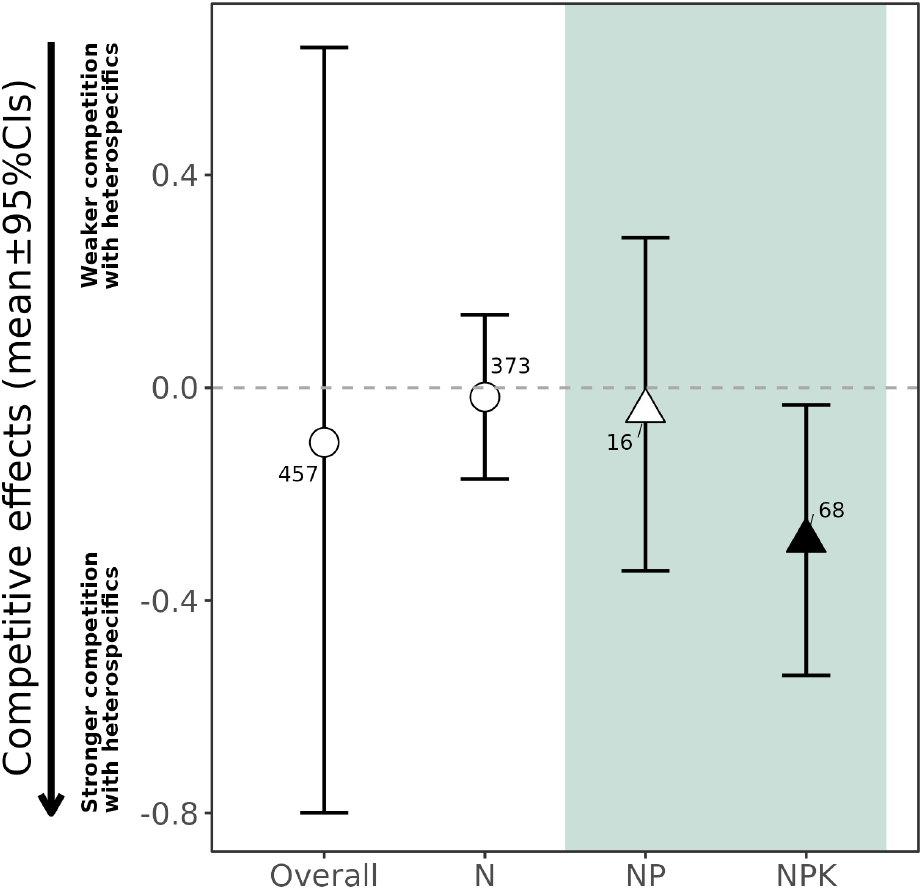
Mean competitive effects with varying nutrient additions as a function of each nutrient type: N, NP, and NPK, with an overall estimate across all types (*n = 457*). Credible confidence intervals that do not cross zero are statistically significant (black filled symbols). Numbers close to mean effects indicate the number of observations. Y-axis: Competition is weaker (positive values) or stronger (negative values) with heterospecifics relative to conspecifics. Overall, nutrient addition tends to intensify competition, with NPK significantly increasing focal plant biomass when grown with conspecifics (i.e., stronger competition in monoculture) compared to heterospecifics.

**Fig 2:**
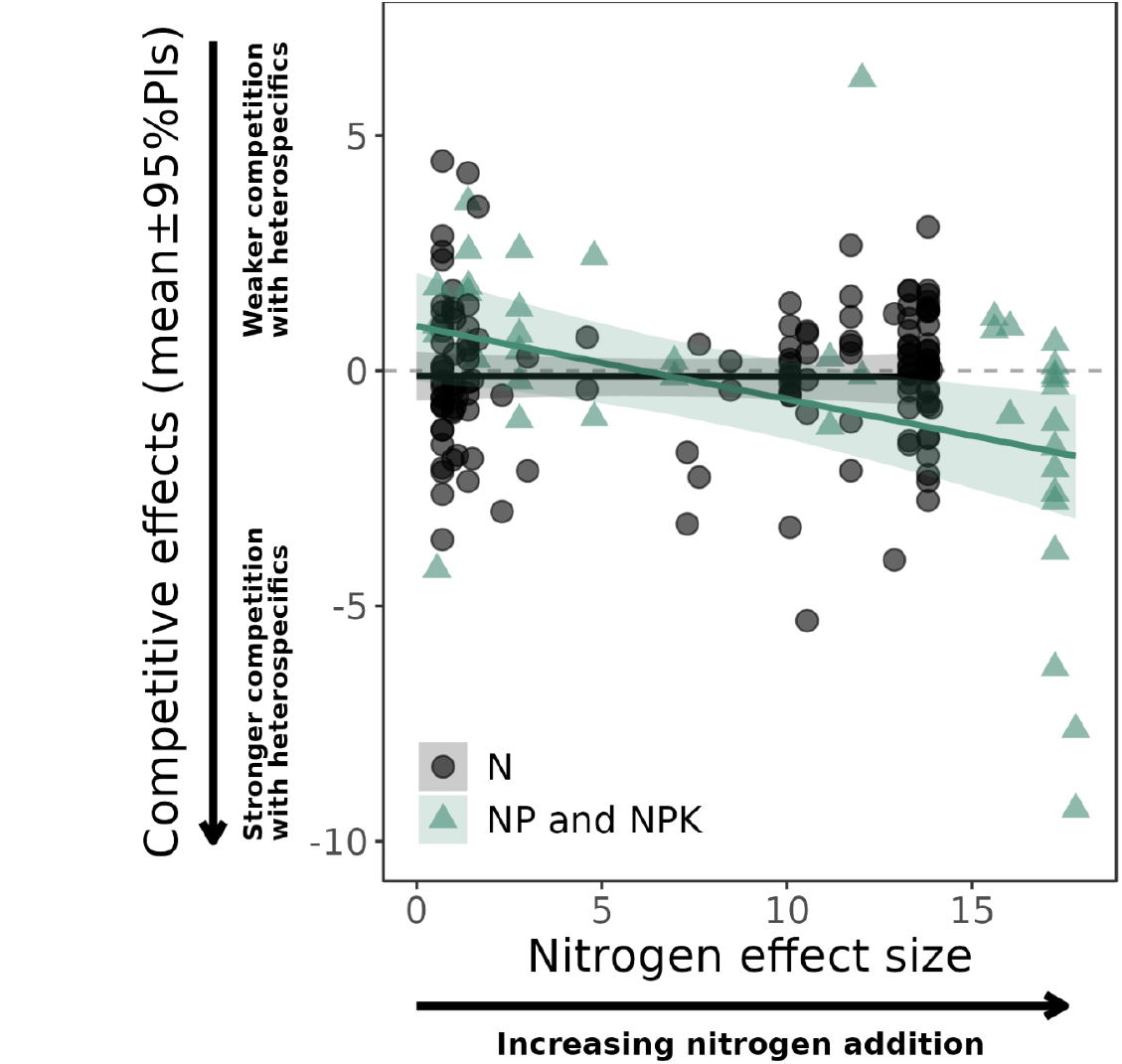
Fold change in competitive effects (ΔLRR i.e., [log response ratio of mixture/monoculture] at high over the low nutrient level) as a function of the magnitude of nutrient addition (i.e., log response ratio of nitrogen quantity at highest over lowest nutrient levels) per nutrient type: N (black) and NP and NPK (green) (*n = 186*). Points are estimated compound competitive effects for each observation. Y-axis: Competition is weaker (positive values) or stronger (negative values) with heterospecifics relative to conspecifics. Solid lines are predicted mean compound competitive effects per category with associated 95% predicted interval (PIs) as shaded areas. Competitive effects are unaffected by N quantity when N is added alone. However, under NP and NPK addition treatments, competitive effects are significantly stronger with heterospecifics relative to conspecifics with each unit change in N added.

**Fig 3:**
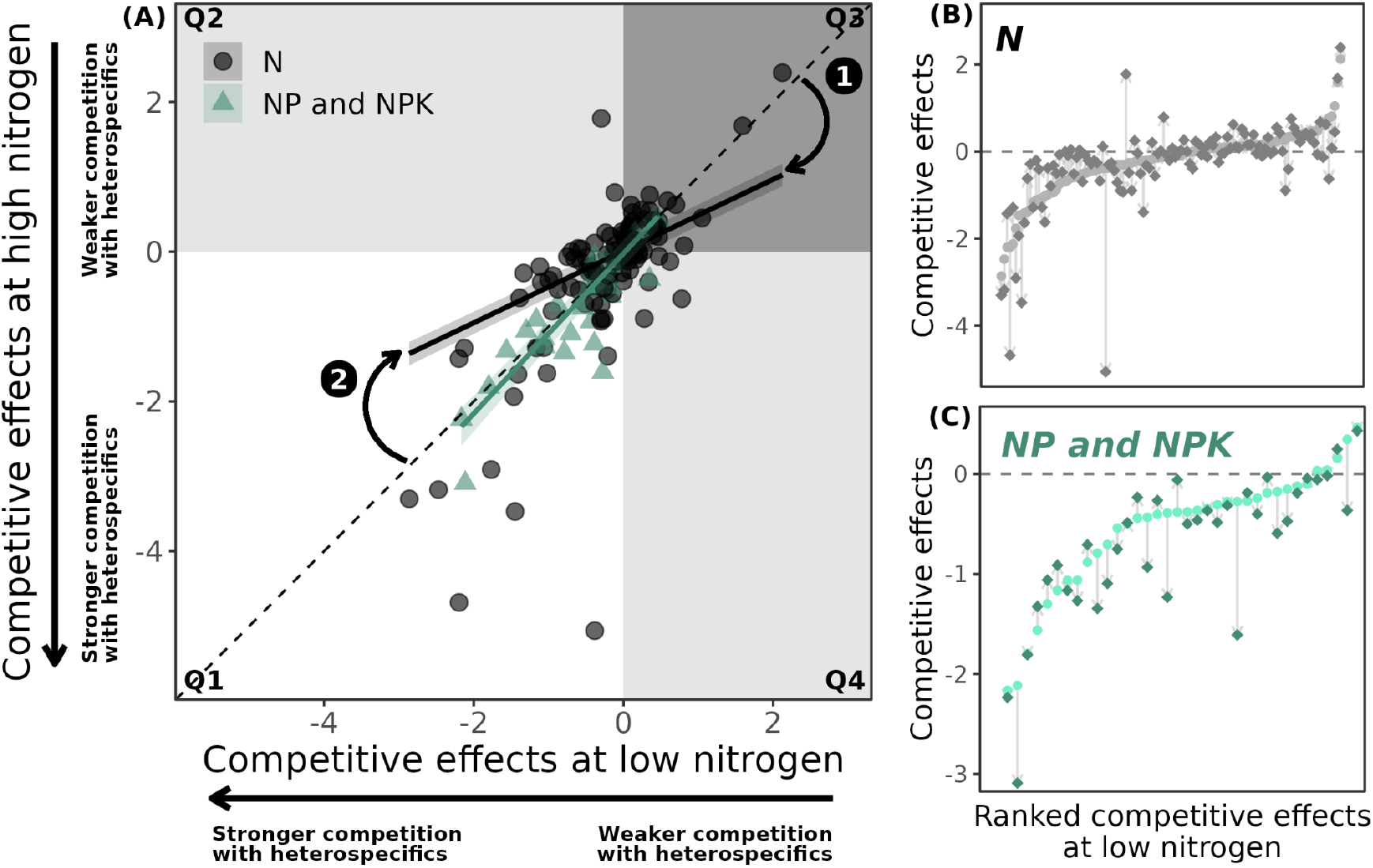
(A) Model results of competitive effects (CE) at highest nitrogen level as a function of CE at the lowest nitrogen level, per nutrient type: N (black), and NP and NPK (green) (*n = 154*). Points are calculated CE for each observation. Solid lines are predicted mean CE per category with associated 95% predicted interval as shaded areas. The four shaded quadrants represent potential competitive outcomes. Quadrant 1 (Q1) (white): Stronger CE with heterospecifics under both low and high nutrients, higher biomass accumulation with conspecifics regardless of nitrogen level. Q2 (light gray): Stronger CE with heterospecifics at low nutrients, but weaker CE with nutrient addition. Q3 (dark gray): Weaker CE with heterospecifics at both nutrient levels, and higher biomass accumulation with heterospecifics regardless of nitrogen level. Q4 (gray): Weaker CE with heterospecifics at low nutrients, but stronger CE with heterospecifics with nutrient addition. The dashed 1:1 line indicates no change in CE between low and high nutrient treatments. Points above or below this line represent a change in CE due to nutrient addition (B, C). Side panels show CI values at low (●) and high (♦) nitrogen levels for N (B) and N-mix (C) treatments, ranked by CI under low nitrogen. Thin, faded arrows connect data points from CI at low to high nitrogen, and larger arrows indicate the overall direction and magnitude of competitive effects shifts for N (B) and NP and NPK (C) treatments. For both axes: Competition is weaker (positive values) or stronger (negative values) with heterospecifics relative to conspecifics. Overall, the effect of nitrogen addition depended on the initial competitive effect under low nutrients. When species experienced weaker competition with heterospecifics at low N, nitrogen addition led to higher biomass accumulation with conspecifics than with heterospecifics. In contrast, when species experienced stronger competition with heterospecifics at low N, nitrogen addition led to higher biomass accumulation with heterospecifics than with conspecifics.

**Fig 4:**
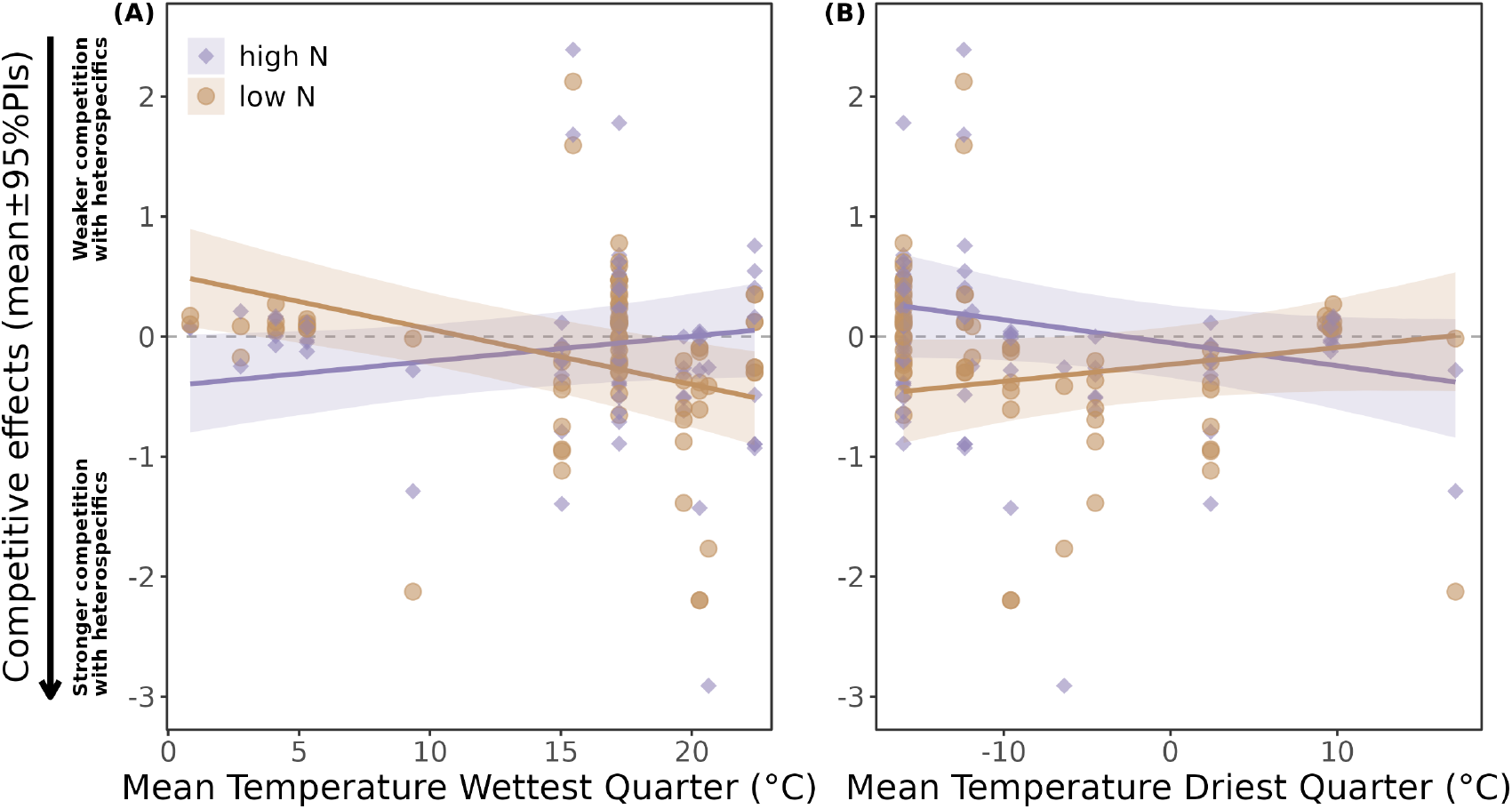
Model results of competitive effects at highest (purple) and lowest (orange) nitrogen levels as a function of the wettest (A) and driest (B) quarters of the year. Points are calculated competitive effects for each observation (*n = 158*). Solid lines are predicted mean competitive effects per category with associated 95% predicted interval (PIs) as shaded areas. The effect of nitrogen addition on competitive effects has interacting outcomes with climate: in the wettest quarter, higher temperatures lead to lower competitive effects values at the lowest nutrient addition levels, reducing biomass accumulation with heterospecifics relative to conspecifics.

### Box: Measuring competitive effects and their interpretations

A species’ competitive ability refers to its ability to exclude other species from a community, whereas niche differentiation, a signature of coexistence, can aid a species’ permanence within a community (14, 30). A crucial step towards understanding species’ competitive ability relies on quantifying <u>**competitive effects**</u>. Competitive effects can be measured as the reduction in performance (i.e., reproduction outcomes, biomass, yield, relative growth rate) resulting from the per capita effects of the presence of heterospecifics vs. conspecifics (12, 13). Here, we illustrate how we leveraged the effects of conspecifics and heterospecifics on focal plant biomass outcomes, a commonly used metric of performance (31), to understand how nutrient addition modifies competitive effects (Box Fig. 1). We quantified competitive effects by calculating the Log Response Ratio (LRR) to plant biomass of focal plants growing with conspecifics and growing with heterospecifics.

#### Visualizing individual variation in species’ competitive effects

To evaluate how individual species’ competitive effects respond to nutrient addition, we divided their competitive effects into two categories: low-nutrient (●) and high-nutrient (♦) conditions (Box Fig. 2A). These are then represented in a two-dimensional space—competitive effects in low-nutrient on the x-axis and high-nutrient on the y-axis. This visualization allows us to track how nutrient addition alters individual variation in competitive effects across species. The dashed 1:1 line represents no change in competitive effects when nutrients are added, while the four quadrants represent distinct competitive outcomes. Side panels show competitive effects ranked by competitive effects under low nutrients. Thin, gray arrows indicate species-level shifts, while means (black dots) and 95% confidence intervals summarize the overall direction of mean response. In Cases I-III, we illustrate the importance of quantifying individual variation: although the mean response is consistently close to zero, it masks substantial individual-level variation in response to nutrient addition

- **Case I:** Nutrient addition shifts species from accumulating higher biomass with conspecifics at low nutrients to higher biomass with heterospecifics at high nutrients.
- **Case II (1:1 line)**: Nutrient addition leads some species to accumulate higher biomass with heterospecifics and others with conspecifics, creating scatter around the 1:1 line (i.e., no consistent directional shift).
- **Case III:** Nutrient addition shifts species from higher biomass with heterospecifics at low nutrients to higher biomass with conspecifics at high nutrients. Beyond these, varying degrees and directions of departure from the 1:1 line reflect other nuanced outcomes, found in Text S3.

## Methods

### Preregistration and Accessibility

The framework, workflow, and protocol for this study were preregistered on the Open Science Framework (32).

#### Systematic search and data extraction

To examine how nutrient addition alters competitive effects, we performed a comprehensive literature search in the Web of Science Core Collection in February 2024, including studies from 1900 to 2023. Our search combined terms related to plant competition and nutrient addition, with both inclusion and exclusion criteria. We used the following Boolean structure:

> *(“plant communit*” OR “plant*” OR “plant population*” OR “species mixture*” OR “pairwise*”) AND (“N add*” OR “N fertiliz*” OR “N suppl*” OR “N enrich*” OR “N gradient*” OR “N level*” OR “P add*” OR “P fertiliz*” OR “P suppl*” OR “P enrich*” OR “P gradient*” OR “P level*” OR “K add*” OR “K fertiliz*” OR “K suppl*” OR “K enrich*” OR “K gradient*” OR “K level*” OR “NP add*” OR “NP fertiliz*” OR “NP suppl*” OR “NP enrich*” OR “NP gradient*” OR “NP level*” OR “NPK add*” OR “NPK fertiliz*” OR “NPK suppl*” OR “NPK enrich*” OR “NPK gradient*” OR “NPK level*” OR “KNP add*” OR “KNP level*” OR “KNP fertiliz*” OR “KNP suppl*” OR “KNP enrich*” OR “KNP gradient*” OR “nutrient* add*” OR “nutrient* fertiliz*” OR “nutrient* suppl*” OR “nutrient* enrich*” OR “nutrient* gradient*” OR “nitr* add*” OR “nitr* gradient*” OR “nitr* level*” OR “nitr* suppl” OR “phosph* add*” OR “phosph* fertiliz*” OR “phosph* suppl” OR “phosph* level*” OR “potassium add*” OR “potassium fertiliz*” OR “potassium suppl*” OR “potassium level*” OR “productivity gradient*”) AND (“biomass*”) NOT (“crop*” OR “AMF” OR “ECM” OR “arbuscular mycorrhizal fung*” OR “ectomycorrhizal fung*” OR “mycorrhiz*” OR “soil respiration” OR “bacterial community” OR “fungal community” OR “cultivar*” OR “plantation*” OR “pasture*” OR “paleo” OR “medicinal” OR “microb*” OR “wheat” OR “pea*”)*.

From our initial pool of results, we selected published peer-reviewed articles according to the following criteria: (1) articles had to report biomass data for a focal plant species grown either in monoculture (i.e., more than one conspecific) or alone (i.e., a single individual), as well as in mixture (i.e., with heterospecifics); (2) plants had to be grown under at least two different levels of nutrient addition and the article had to report the quantity of added nutrient; (3) nutrients added had to include nitrogen (N), phosphorus (P) or potassium (K) applied individually or in any combination. We subsequently excluded studies applying P alone (three studies) since it was biased only towards one growth form, and no study included K-only treatments. We excluded all books, book sections, and conference abstracts. As our focus was on non-agricultural plants, we excluded articles with the following focal plant species: algae, crop (e.g., bokchoy, chipilín, green beans, lettuce, maize, peas, peppermint, rice, soybean, sunflower, wheat), or medicinal plants. We also excluded articles on bioremediation, phytoremediation, phytoextraction, metal uptake, and absorption. We retained articles that included other global change treatments (e.g., elevated CO_2_ or altered water availability) only if they employed a full factorial design (i.e., when all possible combinations of treatments are tested), allowing us to isolate the effects of nutrient addition. We also retained articles when authors clearly identified treatment conditions representing natural or ambient environments.

For each article, we extracted data on the mean and associated variance (standard deviation, standard error, or upper limit of a confidence interval) of aboveground biomass for focal species to calculate our response variable, the log response ratio (LRR) described below. We also recorded data on nutrient type and quantity added at each addition level; the identity, nativity, and growth form of focal and heterospecific species; experiment type (e.g., field experiment, mesocosm, greenhouse); experiment duration; and geographic and climatic information (most commonly, mean annual temperature and precipitation), among other variables (see the complete list in Table S1). We extracted multiple data entries per article since our inclusion criteria required at least two levels of nutrient addition, with each row corresponding to a different nutrient level. Therefore, to account for such non-independence, we added a random effect per study in all models (see ‘Model description and additional analysis’ below).

We extracted data from text, tables, graphs, supporting information, or associated data repositories when available. When data were presented only in graphical form, we used the WebPlotDigitizer online application (33) to extract means and associated variances. For studies that did not report raw data through any of these means, we contacted the corresponding authors directly. For field experiments or mesocosms, we obtained bioclimatic normals (mean annual temperature of driest and wettest quarter) at 10-km spatial resolution from WorldClim2 (34) using the reported study coordinates. We downloaded the bioclimatic normals using ‘geodata’ package (35). To assess the accuracy of this approach, we compared the bioclimatic data with climate values reported in the articles, when available (Fig. S1). In our analysis, we used the WorldClim2 data associated with the location of each study.

We synthetically describe the selection process for the final subset of papers included in this meta-analysis in Fig. S2 as a PRISMA flow chart (36), and provide detailed inclusion and exclusion criteria in Table S2, following the PRISMA Eco-Evo v1.0 guidelines (37). We registered this meta-analysis as a Generalized Systematic Review Registration on the Open Science Framework (32).

#### Data validation

We collaboratively screened and extracted data from studies that met all inclusion criteria, with four recorders (AMB, AR, ADT, and LP) leading this process. During the screening phase, we conducted a trial review of 20 articles, which all recorders read independently. We discussed any inconsistencies to align our interpretation of the criteria and resolve ambiguous cases.

In the data extraction phase, we predefined all variables to be extracted and created a standardized Google Sheets template with drop-down menus to ensure consistent spelling and categorization. We then developed a custom R workflow, ‘collaboratoR’ (38), to interface between collaborative data entry and version-controlled data management using Git. This workflow ingests, validates, and transforms Google Sheets data into CSV files.

We included automated checks for data type consistency (e.g., numeric vs. character), value ranges, and rule-based constraints in the validation process. Once a dataset passed validation, the pipeline pushed it to a GitHub repository, allowing us to track all subsequent changes. Our workflow follows FAIR data principles and aligns with best practices recommended by the Environmental Data Initiative (EDI), including tiered levels for harmonized and derived data products.

#### Effect size calculations

To quantify plant competitive effects, we used the LRR, calculated as the natural log of the ratio of the mean aboveground biomass of the focal species when grown with heterospecifics 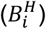 to its biomass when grown with conspecifics 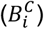 (Box Fig.1). This is a standard approach in the competition literature (e.g., (15, 16, 39). When data of a species grown with conspecifics were unavailable, we used the biomass from available alone-grown treatments as the denominator in the LRR calculation. A small constant (10^−5^) was added to all biomass values to accommodate rare cases where biomass was zero (*n = 8* or 1.75% of the data; (40), either with heterospecifics or conspecifics, due to failure in plant establishment. This adjustment ensured that LRR could be calculated consistently across all studies and species. For each observation *i* within a nutrient level (equation 1):

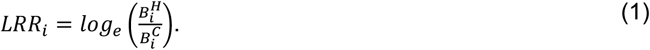

We calculated the pooled sampling variance 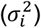 for each observation *i* according to (41), using the sample standard deviations (*S*) and sample sizes (*n*) of the heterospecific (*H*) and conspecific (*C*) groups as:

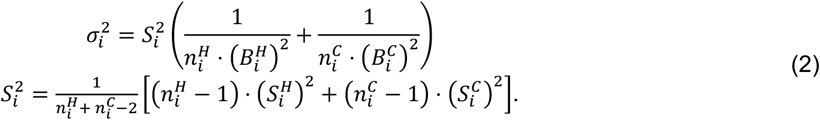

We converted all variance metrics (e.g., standard errors, upper confidence limits) to standard deviations before applying this formula. We used LRR values as the primary measure of competitive effects and used them across analyses addressing Q1 (partially), Q2, and Q3.

To capture the effect of nutrient addition on biomass accumulation, we calculated the change in competitive effects (ΔLRR) as the difference in LRR (equation 1, calculated for each nutrient level) between a high and a low nutrient level. Because nutrient quantities were provided as discrete values along a continuous gradient of nutrient addition across studies, using the high–low comparison allowed us to capture both the magnitude of nutrient addition and its corresponding effect on competitive outcomes. For cases with three or more nutrient treatments, comparisons were performed between successive nutrient levels, for each nutrient type within each study × species combination. We computed ΔLRR within the model framework to enable estimation of missing variances (see below).

To estimate the magnitude of the amount of nutrient added, we calculated the LRR of nutrient addition (Nitrogen effect size; *nutrientLRR*) as the log ratio of the high 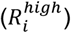 to the low 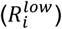 (see paragraph above, same definition for high and low nutrient level comparison) quantity of nutrient applied to a focal species when grown alone, with conspecifics, or with heterospecifics at a given experiment within each study as:

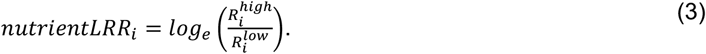

We used ΔLRR, derived from LRR, as the response variable and *nutrientLRR*_!_ as the predictor in a model to partially address question 1 (Q1), as described in the next section

#### Model description and additional analysis

##### Q1: How do nutrients shape plant competitive effects?

To assess how nutrients shape plant competitive effects, we ran two different analyses. First, we fit a model of the calculated LRR as a function of nutrient type (i.e., N, NP, and NPK), including a random effect per study. In this model, we estimated nutrient type intercepts from an overall hyperparameter, *untrt*∼*Normal*(0,1). This analysis focuses on how adding nutrients changes competitive effects and is therefore irrespective of the amount of nutrients added to each experiment.

Second, to test whether nutrient amount modified competitive effects, we analysed the change in competitive effects (ΔLRR) as a function of *nutrientLRR*_!_ (both defined above) for each nutrient type. We also included a random effect per study. For subsequent analyses, we pooled nutrient treatments combining nitrogen with phosphorus or potassium (NP, NPK) because their low sample sizes precluded separate analyses.

##### Q2: How does nutrient addition change species variation in competitive effects?

To investigate how nutrient addition shapes species-specific competitive effects, we selected a subset of data by filtering the LRR of biomass responses at the highest and the lowest nutrient level at each experiment, species x study combination. We fit a model of the calculated LRR of a focal species at the highest nutrient level as a function of its calculated LRR at the lowest nutrient level and study as a random effect. In this analysis, we fit both intercept and slope parameters that varied by nutrient type. The slopes indicate how strongly competitive effects shift from the lowest nutrient level treatment (akin to a 1:1 relationship) to when nutrients are added.

##### Q3: How does nitrogen addition interact with climate to shape competitive effects?

Next, using only field or mesocosm experiments, we analyzed the calculated LRR under nitrogen-only addition at the lowest and highest nutrient levels per species pair and study combination. We modeled LRR as a function of a bioclimatic variable (temperature of the wettest or driest quarter), allowing both intercepts and slopes to vary by nutrient level.

#### Missing variances

We used a Bayesian approach to estimate missing variances (18% of the observations) that would otherwise be excluded in traditional approaches (following (42). Specifically, we estimated any missing variance (σ^2^) as latent variables drawn from log-normal distributions with a mean equal to the logarithm of the largest observed variance among all reported observations, *log*(σ^2^) ∼ *Normal*(*log*(*maxVar*),1). We estimated the fixed-effect parameters using moderately informative prior distributions, *Normal*(0,1) or *Normal*(0,10), depending on parameter type (see code for each specific prior). We estimated study-level (*s*) random effects as *RE*^2^_*S* (i)_∼*Normal*(0, σ^2^_*S*(*i*)_), and variances as σ^2^_*S*(*i*)_∼*Uniform*(0,100). We considered overall coefficients or slopes statistically significant if their 95% credible intervals (CI) did not include zero. We interpreted differences between coefficients as statistically significant when their CIs did not overlap.

#### Model implementation

We implemented all mixed-effects linear models in JAGS (43) through R version 4.3.2. (44) using ‘rjags’ (45). We ran three parallel Markov chain Monte Carlo (MCMC) chains and discarded the burn-in after convergence was reached, following 100,000 iterations. We checked for convergence by visually inspecting trace and density plots. We present posterior means, standard deviations, and 95 % credible intervals (CI) of relevant parameters calculated from 5,000 post-convergence relatively independent samples.

We performed an extensive investigation of potential covariates that could help explain residual variability in LRR by exploring the correlations between the calculated LRR, the response variable, and the candidate covariates compiled from the included articles (Table S1). We followed a similar approach to decide which bioclimatic normals to include in the model, choosing variable(s) with the strongest correlations with LRR. Finally, we investigated publication bias by plotting funnel plots as the inverse variance as a function of calculated LRR (Fig. S3) and performing Egger’s regression tests (Table S3). We performed all data processing and plotting in R version 4.3.2 (44), using the ‘tidyverse’ family of packages (46).

## Results

### Q1: How do nutrients shape plant competitive effects?

We quantified competitive effects as the log response ratio (LRR) of a species’ aboveground biomass when grown with grown with heterospecifics (mixture) relative to conspecifics (monoculture) for each nutrient type and level. Positive LRR values indicate higher biomass production of the focal species when in mixture (i.e., weaker competition with heterospecifics), whereas negative values indicate higher biomass in monoculture (i.e., stronger competition with heterospecifics; Box Fig.1). Then, to examine how nutrient quantity modulates these effects, we calculated the change in competitive effects (ΔLRR) as the log ratio of LRR at high versus low nutrient levels for each fold change in nitrogen addition (nitrogen effect size).

Nutrient addition generally intensified competition with heterospecifics (negative mean); however, with high variability across species (Fig. 1). Neither nitrogen (N) nor nitrogen-phosphorus (NP) additions resulted in significantly stronger competitive effects with heterospecifics (i.e., similar competition with heterospecifics as with conspecifics). In contrast, nitrogen-phosphorus-potassium (NPK) addition significantly intensified competition with heterospecifics, leading focal species to accumulate less biomass when grown with heterospecifics than with conspecifics (Fig. 1). Increasing the amount of nitrogen added alone did not alter ΔLRR (Fig. 2), but adding P or PK alongside nitrogen significantly strengthened competition with heterospecifics when compared to conspecifics. Under NP and NPK, each fold increase in nitrogen addition corresponded to a 1.5-fold increase in competitive effects (ΔLRR).

### Q2: How does nutrient addition change species variation in competitive effects?

We evaluated species’ variability in response to nutrient addition by mapping shifts in competitive effects between the lowest and highest nutrient levels for a given study species. The dashed 1:1 reference line denotes no change in competitive effects between nutrient levels. Points that deviate from this line represent species whose competitive effects either increased or decreased from the lowest to the highest nutrient addition. Therefore, the overall mean competitive effect can remain similar across nutrient levels, but the variation among individual species-pairs reveals whether nutrient addition leads to minimal change or fundamentally tips competitive effects, translating into better performance with conspecifics or heterospecifics.

#### N alone

The response to nitrogen addition depended on the initial competitive effect a species experienced under low-nutrient conditions. When species had higher biomass with heterospecifics under low-nutrient conditions (positive competitive effect values; i.e., weaker competition with heterospecifics relative to conspecifics, quadrant 3 in Fig. 3), nitrogen addition reduced this advantage, shifting the overall competitive effect values toward zero. This shift indicates that species tend to accumulate more biomass with conspecifics than heterospecifics when nitrogen is added alone, reflecting stronger competition with heterospecifics (arrow 1 in Fig. 3A and 3B). However, when species already had higher biomass with conspecifics under low-nutrient conditions (quadrant 2 in Fig. 3), nitrogen addition increased competitive effects values, pulling their overall magnitude towards zero, weakening competition with heterospecifics (arrow 2 in Fig. 3A and 3B). Thus, the overall competitive effects mean of *N* remains close to zero (Fig. 1) even as nitrogen additions increase (Fig. 2), but the variation in responses to added nutrient increases, reflecting divergent responses.

#### NP or NPK

At low nutrients, most species were subject to stronger competition when grown with heterospecifics. With nutrient addition, most species maintained similar negative competitive effects (i.e., the near 1:1 relationship between low and high competitive effects values in green predicted line; Fig. 3A), indicating higher biomass accumulation with conspecifics (33/38 species in Fig. 3A fall in quadrant Q1 [white quadrant], Fig. 3C). Thus, the addition of P and K together with N reinforces strong competition with heterospecifics, as shown in the negative overall competitive effects mean for NP and NPK (significant for NPK, Fig. 1), and by the strong negative responses to increasing N additions in both NP and NPK treatments (Fig. 2).

### Q3: How does nitrogen addition interact with climate to shape competitive effects?

Using a subset of studies that performed field or mesocosm experiments that added only N (*n* = 158; low sample size prevented us from running the analysis with NP and NPK), we investigated how competitive effects at the highest and lowest nitrogen levels responded to mean temperatures during the wettest and driest quarters of the year.

Wettest-quarter temperatures were significantly associated with stronger competition with heterospecifics under low nutrient addition, with competitive effects decreasing by a 0.96-fold per °C (Fig. 4A). However, under high nutrient addition, wettest-quarter temperatures increased competitive effect values, shifting the mean from negative to positive LRR. This corresponds to a change from stronger to weaker competition with heterospecifics, although the effect was not significant (Fig. 4A). In contrast, driest-quarter temperatures at low nutrient levels were associated (marginally significant) with weaker competition with heterospecifics, with competitive effects increasing by a 1.01-fold per °C (Fig. 4B). At highest nutrient addition, driest-quarter temperatures tended to decrease competitive effect values, shifting the mean from positive to negative LRR. This result suggests an increase in competition with heterospecifics, though the effect was not significant (Fig. 4B).

## Discussion

Excessive nutrient addition has consistently led to plant species loss, with clear patterns of diversity decline (8). However, the mechanisms driving plant competitive effects are rarely examined at a global scale. Our meta-analysis, encompassing 71 plant competition experiments from 31 studies worldwide, reveals that nutrient addition increases competitive effects with heterospecifics. This is reflected in higher biomass accumulation of species when grown with conspecifics (monoculture) compared to heterospecifics (mixture), with strongest effect when multiple limiting nutrients were added together (Fig. 1). Nitrogen alone had little effect, but in combination with other nutrients (NP or NPK), adding nitrogen led to stronger competition with heterospecifics relative to conspecifics (Fig. 2). These findings support the niche dimensionality hypothesis, which predicts that reducing limiting resources, and therefore the number of niches, increases interspecific competition (47). Species’ responses to nitrogen addition depended on the initial competitive effect they experienced under low-nutrient conditions. Nutrient addition increased species biomass accumulation with conspecifics when initial competition was weaker with heterospecifics. But it enhanced biomass accumulation with heterospecifics relative to conspecifics when initial competition was stronger with heterospecifics (Fig. 3). These shifts in competitive effects were further amplified by higher temperatures in dry quarters, suggesting that nutrient addition may exacerbate strong competitive effects under extreme conditions (Fig. 4). Together, these patterns highlight how nutrient enrichment and climate interact to reshape plant competitive dynamics and drive potential species loss.

### Q1: How do nutrients shape plant competitive effects?

Adding multiple limiting nutrients to ecosystems intensifies plant competitive effects with heterospecifics (Fig. 1), in line with the niche dimensionality hypothesis (47), which argues that reducing niche dimensions by adding limiting resources intensifies competitive effects (8, 18). Specifically, each fold increase in nitrogen supply with NPK addition resulted in a 1.5-fold decrease in competitive effects compared to nitrogen alone (Fig. 2). In this context, stronger competitive effects with heterospecifics relative to conspecifics align with observations that adding multiple nutrient types tends to accelerate the loss of diversity (8, 48), likely driven by shifts in competitive strengths under nutrient enrichment. These differences may arise because with each additional nutrient type supplied, niche dimensionality declines and nutrient colimitation weakens, reducing competitive effects as species increasingly compete along fewer resource axes (49, 50). This reduction in niche dimensionality may limit the ability of species to coexist, further favoring competitive exclusion (8, 51).

### Q2: How does nutrient addition change species variation in competitive effects?

Nutrient addition can strengthen or weaken a species’ competitive effect depending on how it competes under low-nutrient conditions (Fig. 3). We evaluated species’ variability in response to nutrient addition, revealing how competition shifts between the lowest and highest nutrient levels for a given study species. We found that nitrogen addition reversed overall competitive outcomes. When competition with heterospecifics was weak at low N, adding nitrogen increased competition strength with heterospecifics compared to with conspecifics (Fig. 3, arrow 1 in Q3). These changes in competitive outcomes support a potential mechanism of diversity loss under nitrogen enrichment, which has been shown to reduce species richness by 35%, though rarely causing complete homogenization (2, 52). However, when low *N* levels already promoted strong competition with heterospecifics vs. conspecifics, adding nitrogen reduced competitive effects with heterospecifics, decreasing species biomass (Fig. 3A, B). In contrast, adding multiple nutrient types (NP or NPK) under low-nutrient conditions resulted in consistently stronger competition with heterospecifics, as indicated by lower biomass accumulation when among heterospecifics relative to conspecifics. (Fig. 3, arrow 1 in quadrant Q3). Therefore, adding multiple nutrient types had limited impact on individual competition outcomes (Fig. 3A, C). Our findings on nitrogen’s impact on competition outcomes were consistent with a field study of grass-forb competition, where nutrient addition reversed competitive rankings (53). Specifically, species that exerted weak competitive effects under low-nutrient conditions became stronger competitors after fertilization, while those that were initially strong competitors became weaker (53). Future studies should investigate whether the addition of multiple nutrient types simply reinforces competition strength without significantly altering species responses.

### Q3: How does nitrogen addition interact with climate to shape competitive effects?

In the highest nutrient addition treatments per focal species and study combination, higher temperatures during the wettest quarter (i.e., stressful conditions) weakened competition with heterospecifics relative to conspecifics. Consequently, focal species shifted from producing more biomass among conspecifics to producing more biomass among heterospecifics (Fig. 4A). This shift suggests niche differentiation under stressful conditions. If excess nutrients combined with higher temperatures impose compound physiological stress (e.g., due to metabolic overload, disrupted stoichiometry, or physiological imbalance, etc.), species may shift to facilitative interactions to cope with dual stressors aligning with the stress gradient hypothesis (54, 55). SGH predicts that as environmental stress increases and limits plant biomass accumulation, species interactions shift from stronger to weaker competitive effects, reducing interspecific competition (27, 56, 57).

In contrast, higher temperatures during the driest quarter intensify competition at the highest nutrient level (Fig. 4B). This deviation from SGH, often called the modified SGH, may arise because strong competition can occur at both ends of the stress gradient (58–60). For example, hydric stress can intensify competition among species for the scarce resource (water). In turn, water limitation constrains the ability of plants to buffer nutrient stress (61). When large nutrient additions are not accompanied by sufficient water available to support photosynthetic activity, species may experience resource mismatch, amplifying competition (62). Such resource mismatch can reduce or even halt weak competitive effects (aka facilitation) with extreme stress. Whether this represents a collapse of facilitation in extreme conditions (63) or a nutrient-driven shift from facilitation to competition, as seen in water-limited ecosystems (58, 60), requires further mechanistic testing (62). Thus, the dual pressures of excess nutrient and extreme climates could increase competitive effects, with potential impacts on species diversity maintenance.

### Future Directions

This comprehensive examination of nutrient-driven mechanisms in explaining species’ competitive effects can be expanded in four key ways. *<u>First</u>*, assessing how nutrient addition shapes coexistence mechanisms will require estimating full per capita interaction coefficient matrices (e.g., the competition coefficients *a*_*i j*_, *a*_*ii*_ *sensu* dynamic population models) within the framework of contemporary niche (64) or modern coexistence theory (30). Our analysis focuses on one side of the interaction: how nutrient addition alters the effects of competition on a focal species, since most existing studies do not use designs like replacement series or minimal surfaces (13) that allow estimation of coefficients to evaluate niche and fitness differences (53). *<u>Second</u>*, our meta-analysis reveals an ecological gap: most experiments focus on N, NP, or NPK additions, and most field studies are restricted to nitrogen addition alone. This focus is reinforced by the bias of the dataset on temperate plant species (Fig. S4), where nitrogen is often the primary limiting nutrient. Such narrow focus limits our ability to test how other niche dimensions influence competition via co-limitation (11). For instance, in alpine meadow communities, nitrogen most strongly limited species competition, with significant effects in all nitrogen-containing treatments. However, the absence of effects under P, K, or PK additions calls into question the consistency of the niche dimensionality hypothesis (65). By contrast, tropical ecosystems are frequently phosphorus-limited, suggesting that temperate-focused findings may not capture the full range of nutrient-driven competition globally (66, 67). *<u>Third</u>*, while synergistic and strict co-limitation are well documented (68), most studies, including ours, rely on factorial designs and broad response categories. Future work should adopt approaches to capture whether nutrient effects are saturating (sub-additive), hierarchical (serial limitation), or dependent on the order of nutrient addition. These insights can help uncover the underlying structure of resource limitation for plant competitive effects (69, 70). *<u>Finally</u>*, while our focus was on plant species from natural communities, the same framework could be applied to crop systems by quantifying how fertilizer alters competitive effects. For instance, high nitrogen inputs can reduce soil organic carbon, suggesting that nutrient imbalances, especially under stress, may destabilize crop systems (71). Tracking competitive effects under high nutrients in dry seasons could improve agricultural outcomes and help protect natural systems, anticipating the effects of global change.

## Conclusion

Nutrient enrichment from agricultural runoff and anthropogenic changes are major global change drivers of species diversity loss. While patterns of diversity loss from nutrient deposition are clear, the processes driving competitive effects are rarely examined on a global scale. Our global meta-analysis reveals that nutrient addition, particularly multiple-nutrient enrichment, intensifies competitive effects favouring monocultures, as indicated by species reducing biomass accumulation when growing with heterospecifics relative to conspecifics. These changes to competitive effects could lead to a decline in diversity. Temperature and water supply interact with nutrient enrichment to amplify these competitive asymmetries, likely further shaping species dominance patterns under climate change. Our findings highlight the urgent need for a global reconsideration of nutrient management strategies to mitigate the unintended consequences of fertilization and nutrient deposition on plant diversity and ecosystem stability.

## Acknowledgements and funding sources

This study was funded by The Institute for Biodiversity, Ecology, Evolution, and Macrosystems (IBEEM) (https://ibeem.msu.edu/) at Michigan State University (PI: PLZ) through the Michigan State University Strategic Partnership Grant Program. IBEEM Postdoctoral Fellowships provided support to AMB, AR, LP, and ADT; AR was additionally supported by the MSU EEB Presidential Postdoctoral Fellowship. This work was supported in part through computational resources and services provided by the Institute for Cyber-Enabled Research at Michigan State University. We thank all authors who generously shared their data.

## Author contributions

LP, AR, AMB, ADT conceived of the study, LP, AR, AMB, ADT designed the study with guidance from LLS and PLZ. LP, AR, AMB, ADT, PB collated and synthesized the data. LP (lead), AR, AMB, and ADT (support) curated the database based on the workflow developed by PB. LP (equal) and AR (equal) performed the analysis and wrote the first draft of the manuscript with input from all co-authors. LLS (equal) and PLZ (equal) supervised the manuscript; PLZ acquired funding. All authors contributed to editing and approve of the final draft.

